# Wnt7a is a Novel Lymphangiocrine Factor Driving Cholangiocyte Proliferation during Liver Regeneration

**DOI:** 10.1101/2025.10.16.682805

**Authors:** Aarti Sharma, Deepika Jakhar, Pinky Juneja, Jayesh Kumar Sevak, Ashwini Vasudewan, Simran Sharma, Impreet Kaur, Veluru Gautham, Viniyendra Pamecha, Dinesh M Tripathi, Shiv K Sarin, Sumeet Pal Singh, Savneet Kaur

## Abstract

The significance of lymphatic vessels (LVs) and lymphatic endothelial cells (LyECs) during liver regeneration remains unexplored. We aimed to characterize contribution of LVs and LyECs in liver regeneration using two-thirds partial hepatectomy (PHx) models. An increased number and diameter of CD31+PDPN+ LVs was observed at day 2 and 5 post-PHx. Proteomic analysis of LyECs and ELISA studies in conditioned media of LyECs revealed that one of the secreted proteins, Wnt7a was uniquely expressed at day 2 post PHx. In vitro studies confirmed that Wnt7a is a potent stimulator of proliferation for both hepatocytes and cholangiocytes. In vivo inhibition of lymphangiogenesis and reduction in Wnt7a led to a decline in the percentage of CK19+ cholangiocytes and bile ducts. Western blot analysis showed that Wnt7a activated the downstream Frizzled 7 receptor and non-canonical planar-cell-polarity pathway to drive the proliferation of cholangiocytes. In vivo, Wnt7a loss-of-function experiments in zebrafish along with inhibitor studies in rat models of PHx further re-iterated clear links between Wnt7a and proliferation of bile ducts. Pro-regenerative effects of Wnt7a were demonstrated in a model of small-for-size syndrome (80% PHx), where administering recombinant Wnt7a enhanced both cholangiocyte and hepatocyte proliferation. Elevated Wnt7a levels were seen in donors of liver transplant patients at day 1 and 2 post-hepatectomy and Wnt7a also facilitated growth of human cholangiocyte organoids in vitro. Our findings uncover the novel role of Wnt7 as one of the key lymphangiocrine signals that facilitates proliferation of cholangiocytes during liver regeneration.

**Highlights:**

- Number of functional lymphatic Vessels (LVs) and lymphangiogenesis is enhanced between day 2 and day 5 after partial hepatectomy (PHx) in rat models and liver transplant donors.
- Wnt7a released from lymphatic endothelial cells (LyECs) acts as a potent lymphangiocrine factor, stimulating proliferation of hepatocytes and cholangiocytes in vitro, across both rodent and human models.
- Inhibition of lymphagiogenesis during PHx transiently reduces number of cholangiocytes during early liver regeneration.
- Wnt7a signals through Frizzled 7 receptor to activate p-AKT, p-JAK1 and p-STAT3 proteins in cholangiocytes.
- Administration of recombinant Wnt7a in 80% PHx models induces proliferation of hepatocytes and bile ducts.

## Introduction

Vessels serve as channels to transport oxygen, nutrients and metabolites to the tissues and also to export waste products^1^. After ischemia during tissue regeneration or injury, endothelial cells that line the vessels in response to tissue injury signals re-establish the capillary network in a coordinated sequence of events to keep the tissue oxygen and nutrient supply intact via the process of angiogenesis^2^,^3^,^4^. Besides acting as conduits for oxygen and metabolite delivery and regulating angiogenesis, recent studies have suggested that endothelial cells and vessels also have key paracrine and/or angiocrine functions during tissue regeneration and disease^5^. Among various organ-specific endothelial cells, the liver sinusoidal endothelial cells (LSECs) significantly contribute to balancing liver regeneration and fibrosis via secretion of a multitude of angiocrine factors^6^. Liver is a highly regenerative organ and signals from the vascular bed or LSECs profoundly control proliferation of hepatocytes to restore liver functions during regeneration. It has been reported that the pericentral LSECs secrete Wnt ligands including Wnt2 and Wnt9 and growth factors like HGF, which play an essential role in hepatocyte proliferation^7^,^8^.

Besides the blood capillaries, liver contains a network of lymphatic capillaries that are essential for fluid homeostasis and immune response regulation^9^,^10^. Tissue fluid in the sinusoids which is rich in proteins and other macromolecules released from the LSECs is drained as lymph in the lymphatic capillaries towards larger lymphatic vessels (LVs) in the portal areas and from their it is drained into the extrahepatic lymph nodes and finally into the thoracic duct^11^. The intrahepatic lymphatic capillaries are not overlapped by pericytes or smooth muscle cells. The endothelial cells of the lymphatic capillaries i.e. lymphatic endothelial cells (LyECs) also contains pores and have specialized button like junctions^9^. There are now several recent evidences that lymphatic vessels and LyECs have other physiological functions other than fluid drainage and regulation of immune cell responses^12^. Lymphatic vessels and LyECs are known to play an important role in regeneration and activation of stem cells in other organs like intestine^13^, heart^14^, brain^15^, hairs and skin^16^. The role of hepatic LyECs in liver growth and regeneration however has remain largely unexplored mainly because of the presence of overlapping markers of these cells and LSECs. In the current study, we hypothesized and investigated the role of hepatic LVs and LyECs during liver regeneration.

## Results

### Lymphangiogenesis occurs two days post PHx

To investigate the contribution of lymphatic vessels (LVs) and lymphatic endothelial cells (LyECs) during liver regeneration, rat models of 70% PHx were prepared and different liver cells were studied at various time points such as 12h, D1, 2, 5, 7 and 15 and compared with their respective shams (**Fig1A**). Liver to body weight ratio showed a significant reduction at 12h (2.8 fold, p<0.0001), D1 (2.78 fold, p<0.0001) and D2 (0.38 fold, p<0.001) compared to sham, after which it began to increase and total liver mass regained at D7 to D15 comparable to sham (**Fig1B**). Immunostaining and flow analysis showed an increased percentage of proliferating PCNA+ hepatocytes at D1 (86.7±10.10%, p<0.001) and D2 (87.5±7.1%, p<0.001) (**SFig1A-C**). CK19+ cholangiocytes were found maximum during D2 as seen by both immunohistochemical (IHC) (50.2±4.1%, p<0.01) staining (**SFig1D&E**) and flow analysis (63.6±5.17%, p<0.0001) (**SFig1F**). Kupffer cells/CD68+ macrophages were also studied and their numbers were increased at D2 post-PHx as shown by flow analysis (45.5±7.65%, p<0.001) (**SFig1K**).

**Figure 1:**
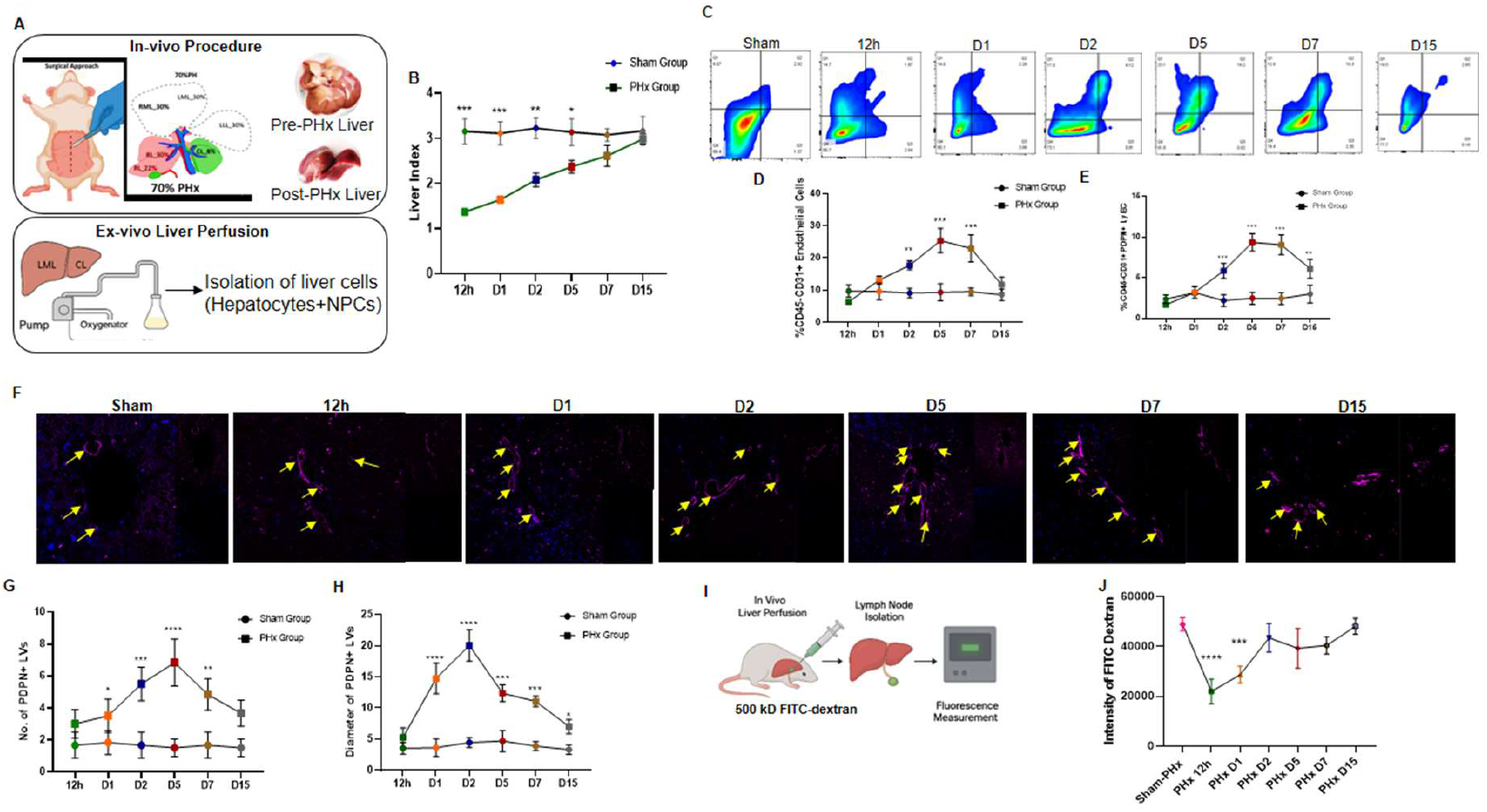
Characterization of Liver Regeneration and Lymphatic Vessel Dynamics Following 70% Partial Hepatectomy (PHx). Diagram illustrating the collagenase perfusion and digestion method used to isolate various liver cell populations from rat models following 70% PHx at various time points from 12h to D15 **(Fig1A)**. Measurement of the liver-to-body weight ratio in rats following 70% PHx, compared to sham-operated controls, at various time points (12h, D1, D2, D5, D7, and D15). There was an initial significant reduction at 12h, D1, and D2, followed by mass regain, reaching near-total regeneration by D7-D15 (**Fig1B**, mean ± SD, n=3, *p <0.05, **p <0.01, ***p<0.001). Results of flow cytometry analysis showing the percentage of CD45−CD31+ and CD45−CD31+PDPN+ endothelial cells (**Fig1C**, mean ± SD, n=3, **p <0.01, ***p<0.001). Graphical representation demonstrating a maximum endothelial population size at Day 5 (D5) post-PHx (**Fig1D**, mean ± SD, n=3, **p <0.01, ***p <0.001). Flow cytometry quantification of CD45−CD31+PDPN+ LyECs showed a maximum no. at D2&D5 (**Fig1E**, n=4, mean ± SD, **p <0.01,***p <0.001). Representative immunofluorescence images of PDPN+ LVs **(Fig1F)** and quantification confirming the flow cytometry findings (**Fig1G**, scale bar=40μm, mean ± SD, n=6, *p <0.05, **p <0.01, ***p <0.001). Quantification of LV diameter (based on PDPN staining) from immunofluorescence data (**Fig1H**, scale bar= 40μm, mean ± SD, n=3, **p <0.01, ***p <0.001). Line plots showing data from the FITC-dextran drainage assays at D2, D5, and D7 post-PHx (**Fig1I&J**, mean ± SD, n=3, ***p <0.001).

Among the liver endothelial cells, maximum number of CD31+ endothelial cells were observed at D5 (66.7±3.8%, p<0.001) after PHx via IHC (**SFig1G&H**) compared to sham. CD31+PDPN+ LyECs maximally at D2 (64.7±6.07%, p<0.0001) & D5 (72.8±5.3%, p<0.0001) post-PHx compared to sham (**Fig1C-E**). Both immunofluorescence and IHC imaging clearly revealed that PDPN+ LVs were decreased at the initial time points of 12h and D1 after PHx and then began to increase at time points of D2, D5, D7 and D15 (**Fig1F&G & SFig1I&J**). The diameter of LVs was also maximally increased at D2 (80±5.59%, p<0.0001) (**Fig1H**). Next, the drainage of FITC-dextran was measured to assess the functionality of LVs (**Fig1I**). The drainage function decreased at the initial time periods i.e. at 12h (58±6.9% p<0.001) and D1 (50±9.1%, p<0.001) but then it increased at D2, 5, 7 and 15, comparable to sham (**Fig1J**), indicating restoration of LV functionality. The number of CD68+ macrophages was also maximum at D2 post-PHx as shown by flow analysis (32±8.1%, p<0.0001). This data clearly indicated that hepatic **lymphangiogenesis and growth of functional LVs occurs between 2-5 days post-PHx**.

### Proteomic and secretome analysis of LyECs identify Wnt7a as one of the protein during early liver regeneration

Next, we aimed to dissect specific protein signatures of LyECs at D2 post-PHx in comparison to sham. For **this, we first ensured the isolation of hepatic LyECs (Lymphatic Endothelial Cells) by isolating the liver vascular cells** from perfused livers as per previous protocols^17^. The vascular cells consisted of **all liver endothelial cells (LECs)**. For further isolation of LyECs, cells were labelled with **CD45, CD31, and PDPN markers** and subjected to **FACS sorting** to isolate LyECs (CD45-CD31+PDPN+) and LECs (CD45-CD31+PDPN-) (**Fig2A***). Expression of LV-specific markers i*.*e. Prox-1*(10.41 fold, p=0.009*), PDPN* (9.16 fold, p<0.0001*), VEGFR-3* (7.5 fold, p=0.0005*) and LyVE-1* (6.67 fold, p=0.0004*) was upregulated in LyECs in comparison to LECs at D5 post PHx, confirming lymphatic identity of the cells (****Fig2B***). After confirming LyEC identity and over 90% purity and viability (via trypan blue), these sorted cell populations were used for **proteomic analysis**. We employed a **Fast-seq approach** for proteome identification. **Biological triplicates** of LyECs (Sham PHx LyECs and PHx D2 LyECs) yielded a total of approximately 6600 protein identifications. Of these, 227 were **up regulated** and 84 **down regulated**, as shown by volcano plots (log2FC > 1.5, p < 0.05, **Fig2C**). **Principal Component Analysis (PCA)** revealed **inter-group variations** of 68.6% and **intra-group variations** of 3.35% between PHxD2 and Sham PHx LyEC (**SFig2A**). **Unsupervised hierarchical clustering** of these genes was performed, and the resulting heat map distinctly segregated individual sample abundance into different clusters for LyECs (**SFig2B**). We performed **pathway enrichment analysis** on differentially expressed genes (DEGs) to elucidate KEGG pathways. In **PHx vs Sham LyECs**, KEGG analysis showed **upregulation** of major pathways such as HIF-1 signalling, cAMP signalling and regulation of actin cytoskeleton. **Downregulated** pathways included Rap1 signalling, insulin secretion etc (**Fig2D**). Gene ontology analysis showed the 72 upregulated and 41 downregulated biological processes (BP) terms, the cellular component terms (CC) were 56 upregulated and 33 downregulated, the molecular function (MF) terms were 62 upegulated and 38 downregulated (**SFig2C&D**).

**Figure 2:**
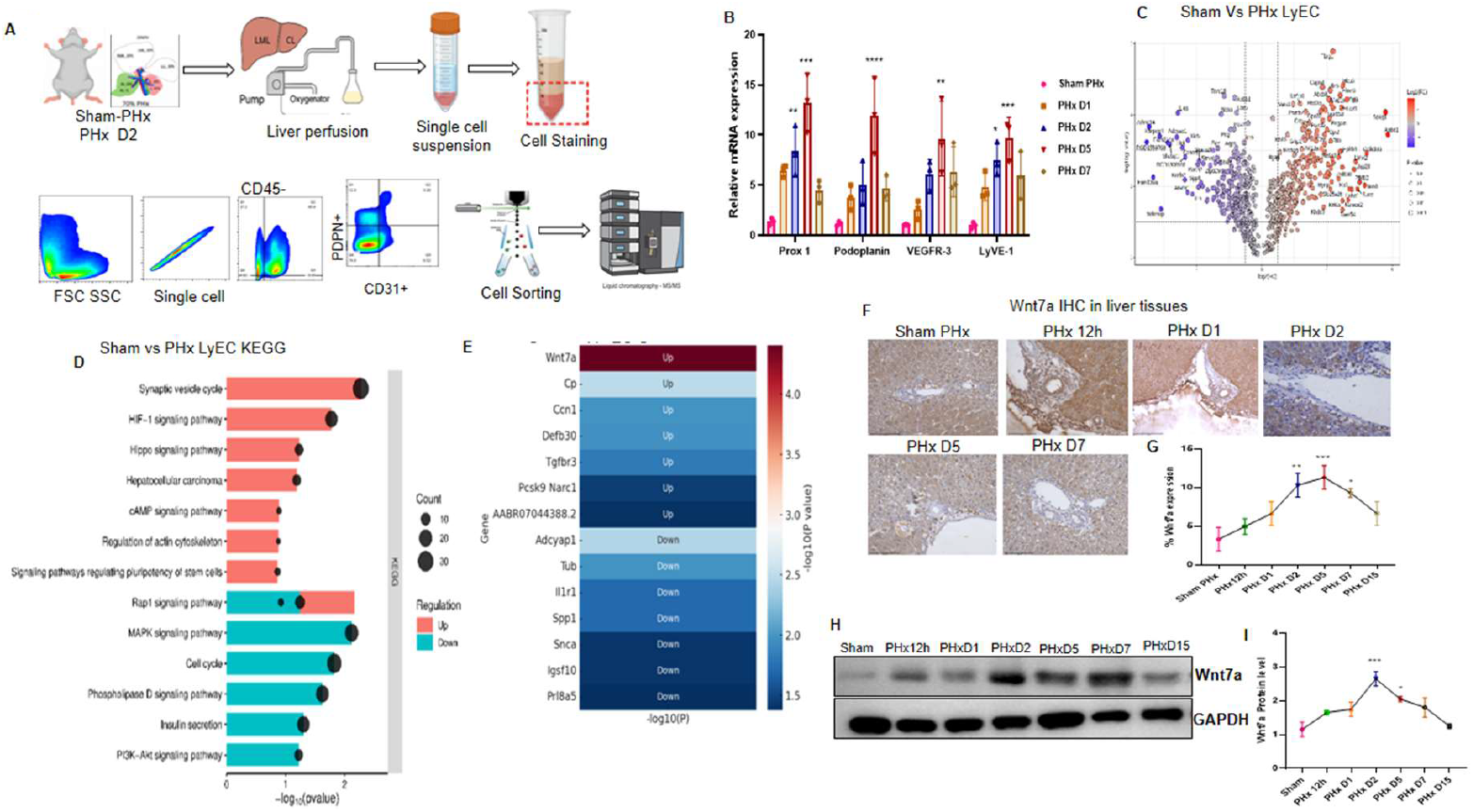
Isolation and Proteomics Profiling of Liver Lymphatic Endothelial Cells (LyECs) following Partial Hepatectomy (PHx). Representative flow cytometry plots showing the gating strategy used to isolate liver endothelial cells (LECs, CD45−CD31+) and subsequent sorting of LyECs (CD45−CD31+PDPN+) from the total endothelial fraction of rat livers 2 days (D2) post-PHx (**Fig2A**). Real-time PCR (RT-PCR) analysis showing the up-regulation of key lymphatic markers (Prox1, Podoplanin, VEGFR−3, and LyVE−1) in sorted LyECs at D2 and D5 following PHx compared to Sham. Error bars represent mean±SD (**Fig2B**, n=3, **p <0.01, ***p <0.001, ****p <0.0001). Gene expression using volcano plots comparing LyECs from PHx D2 animals versus Sham animals (**Fig2C**). Kyoto Encyclopedia of Genes and Genomes (KEGG) enrichment analysis showing major biological pathways that are significantly up-regulated and down-regulated in LyECs at PHx D2 compared to sham LyECs (**Fig2D**). Clustering analysis visualized by a heatmap showing the expression profile of top up-regulated and down-regulated secretory genes in LyECs from PHx D2 compared to Sham (**Fig2E**). Representative IHC staining (**Fig2F**) and quantification (**Fig2G**) for Wnt7a in liver tissues harvested from 12h to D15 post-PHx. Scale bars: 200μm (n=3, *p <0.01, **p <0.01, ***p <0.001). Representative protein level (**Fig2H**) and quantification (**Fig2I**) confirming the time-dependent increase in Wnt7a protein expression at D2 post-PHx. GAPDH was used as control.

Next to study angiocrine factors, we enlisted differentially expressed secretory factors in LyECs during PHx. We categorized the proteins using UniProt to identify secreted proteins among the significant genes. **Results** illustrated upregulated DEGs like **Defb30, Ccn1, Pcsk9, Tgfbr3, Cp, Wnt7a** (**Fig2E**) and downregulated DEGs such as **Snca, Prl8a5, Tub, Il1r1, Igsf10, Spp1, Adcyap1** (**Fig2E**) in LyECs. Based on its involvement in KEGG pathway and gene ontology analysis such as PI3K, MAPK, signalling pathways related to pluripotency of stem cells, hepatobiliary cells proliferation etc, of all the secreted proteins, we selected one of the uniquely expressed gene i.e. Wnt7a, a growth factor of significant importance and dissected its role in further studies. IHC showed an increased hepatic expression of Wnt7a in periportal areas at D2 post-PHx compared to sham (**Fig2F**). A kinetic analysis revealed that its expression starts increasing from 12h till D5 by about 3 fold **(p <0.0001**) after which, it showed a decreasing trend (**Fig2G**). Given the fact that Wnt7a is a secreted protein, next, we studied its expression in the secretome of LyEC cultures. However, given the very small percentage of hepatic LyECs, we were not able to successfully culture these cells alone. To address this, we sorted and cultured the entire CD31+ fraction and collected their secretome for western blot analysis at different time points. There was an increased expression of Wnt7a at D2 (**2.33 fold, p =0.0008**) post-PHx compared to sham (**Fig2H&I**). **This concluded that Wnt7a is expressed in the secretome of liver vascular cells and its expression is highest at day 2 post-PHx**.

### Wnt7a from LyECs enhances proliferation of both hepatocytes and cholangiocytes

Next, we investigated the specific expression of Wnt7a in the LyECs, including, CD31+PDPN+ cells and other CD31+PDPN-vascular cells in PHx (**Fig3A**). In the existing database of human liver single cell RNA sequencing data^18^, both LSECs and LyECs were shown to be a source of Wnt7a in human liver (**SFig3A&B**)^18^. In our analysis, at D2 post-PHx, the %Wnt7a+ CD31+PDPN+ were 30±8% (p=0.008) more as compared to %Wnt7a+ CD31+PDPN-(**Fig3B**). Gene expression analysis also showed that Wnt7a expression was significantly up-regulated at D2 (2.28 fold, p<0.001) post-PHx in CD31+PDPN+ cells compared to CD31+PDPN-cells (**Fig3C&D**). Having established that Wnt7a is a secreted protein and is majorly expressed by CD31+PDPN+ LyECs, we checked its angiocrine effect on both hepatocytes and cholangiocytes. We first tested the effect of LyEC-CM treatment on hepatocytes and cholangiocytes organoids in 3D cultures. There was an increased growth of hepatocyte organoids (with diameter 1.56 fold, p= 0.025) (**SFig3C&D**) and cholangiocytes organoids (diameter increased by 2.23 fold, p=0.006 (**SFig3E&F**). Next, we tested the effect of rWnt7a on hepatocytes and cholangiocytes. A substantial increase in number (55.2±7.6%, p=0.0024) and diameter (1.83 fold, p=0.0349) of PCNA+ cholangiocytes was observed in the presence of rWnt7a **(Fig3 E-G**). We also observed an increased number (37.5±8.3%, p=0.04) and diameter (1.5 fold, p=0.03) of PCNA+ hepatocytes in presence of Wnt7a, although this increase was less than that observed in cholangiocytes (**Fig3H-J**). In fact, in presence of rWnt7a, hepatocyte organoids showed an increased expression of biliary progenitor markers including ov-6 (58.5±3.02%, p=0.003) as shown by imaging and flow analysis (**Fig3K and SFig3G&H**). This implied that Wnt7a released from LyECs might be involved in inducing the proliferation of cholangiocytes. We also co-cultured LyECs and cholangiocytes together and observed an increased organoid growth (48.5+5.6%, P=0.001) compared to controls (**Fig3L&M**). **These results suggested that Wnt7a secreted from LyECs facilitated the proliferation of both hepatocytes and cholangiocytes in vitro**.

**Figure 3:**
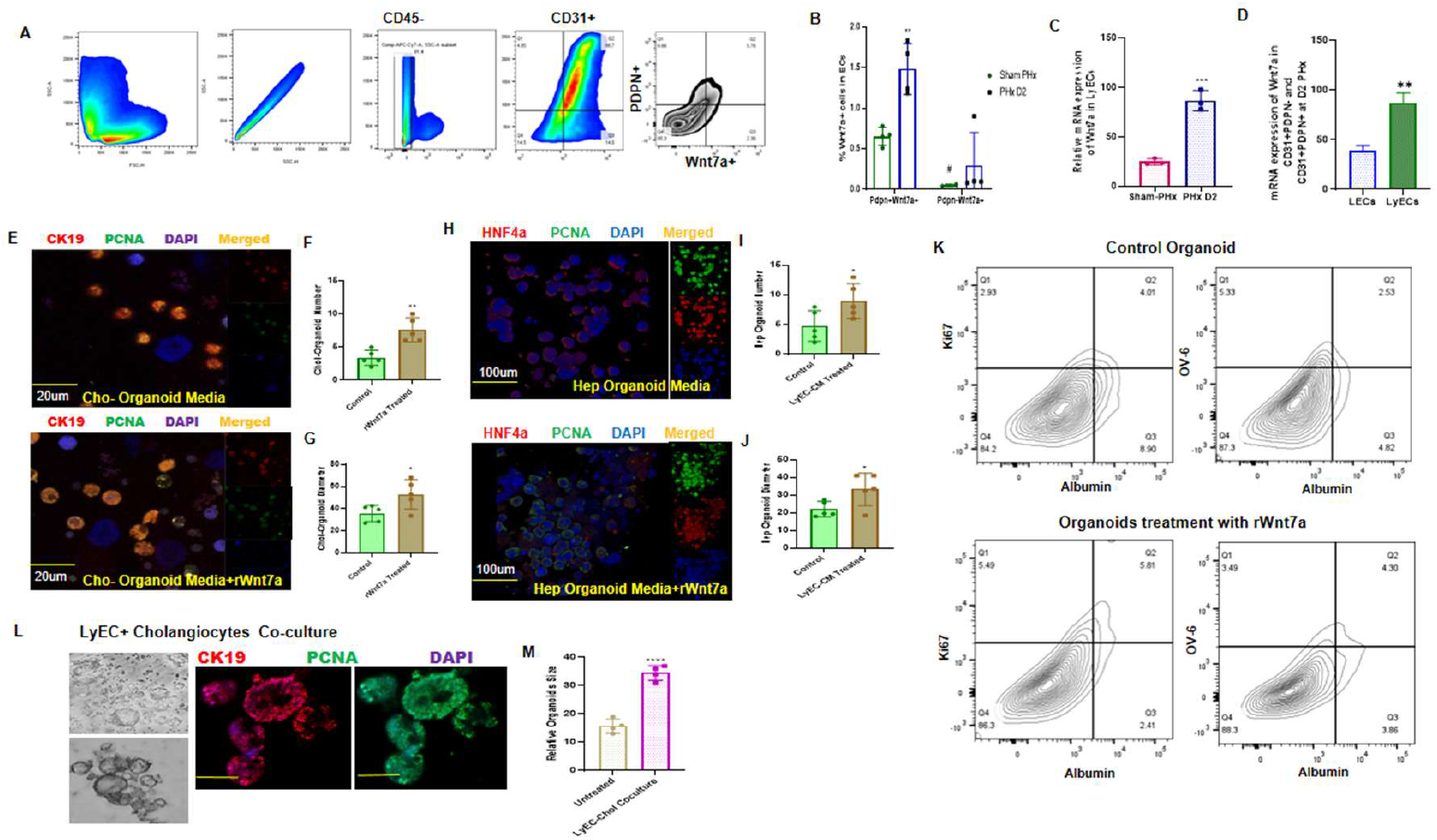
Wnt7a Secreted by LyECs promotes the proliferation of hepatocytes and cholangiocytes in 2D and 3D Cultures. Representative flow cytometry plots for Wnt7a expression in endothelial subpopulations post-PHx (**Fig3A**) and quantification showing the percentage of Wnt7a+ cells within the total endothelial fraction (CD31+) and lymphatic endothelial fraction (CD31+PDPN+) (**Fig3B**, n=3, mean ± SD, n=3, *p <0.05, **p <0.01). RT-PCR analysis in sorted hepatic LyECs from sham and D2 compared to both Sham LyECs and PHx LECs. Error bars represent mean ± SD (**Fig3C&D**, n=3, **p <0.001, ***p <0.0001). Representative immunofluorescence images, quantification of the number of PCNA+ cholangiocytes, and quantification of cholangiocyte diameter in 3D culture following treatment with recombinant Wnt7a (rWnt7a) (**Fig3E-G, n=3, *p <0.05, **p <0.01**). Representative immunofluorescence images, quantification of the number of PCNA+ hepatocytes, and quantification of hepatocyte diameter in 3D culture following rWnt7a treatment (**Fig3H-J**, n=3, *p <0.05, **p <0.01). Flow cytometry analysis showing that hepatocytes cultured in the presence of rWnt7a begin to express biliary markers (**Fig3K**, *p <0.05). Immunofluorescence imaging and quantification of cholangiocyte organoids directly cultured on a monolayer of LyECs (**Fig3L&M**, *p <0.05).

### Inhibition of lymphangiogenesis and reduced expression of Wnt7a leads to a decrease in cholangiocyte proliferation

We next probed if inhibition of lymphangiogenesis and a reduction in LyECs affects Wnt7a levels and liver regeneration in PHx models. For this, 70% PHx rats were given MAZ51 (a VEGFR-3 inhibitor) intraperitoneally during surgery and they were compared with their respective shams at different time points (**Fig4A**). The CD31+PDPN+ LVs were significantly decreased at D2 (0.6 fold, p <0.0301) in the MAZ51 group as compared to untreated PHx group (**Fig4B**). The CD31+PDPN-endothelial cells also showed a significant reduction at D2 (40±4.19%, p <0.03) & D5 (35.7±7.2%, p <0.03) in the treated groups compared to untreated ones (**SFig4A**). Importantly, Wnt7a protein expression showed a substantial decrease in MAZ51 treated animals at D1 (53.1±4.6%, p <0.001) and D2 (57.1±9.2%, p <0.001) post-PHx in comparison to untreated PHx group as seen by both western blotting and further Wnt7a expression regained at later time points i.e. D5 & D7 (**Fig4C&D**). HE and MT data showed no significant damage or injury in the liver due to MAZ51 treatment at any time post PHx (**SFig4B&C**). In terms of liver growth, there was however only a minor and insignificant decrease in liver index at D2 (27.7±1.3%, p <0.12) post-PHx in MAZ51 treated group as compared to that seen in untreated groups. There was also no change in liver index at other time points in MAZ51 treated group versus untreated ones (**Fig4E**).The percentage of PCNA+ hepatocytes also did not change after MAZ51 treatment (**SFig4D&E**). MAZ51 treatment however substantially decreased the CK19+ cholangiocytes at D2 (33.3±5.5%, p <0.03) and D5 (35.3±7.2%, p <0.007) post-PHx (**Fig4F)**. The number of macrophages also significantly reduced at D2 (35.7±6.3%, p <0.03) in MAZ51 treated animals (**SFig4F**). Since there was a major effect on cholangiocytes, next, we compared the kinetic analysis of both PDPN+ LVs & CK19+ bile ducts **(BD)** in the liver tissues. We observed a similar kinetic analysis between these two cells at different time-points in PHx (**Fig4G&H**) and MAZ51 treated PHx groups (**SFig4G&H**). This indicated that Wnt7a might be one of the key lymphangiogenic factor supporting cholangiocyte proliferation in vivo.

**Figure 4:**
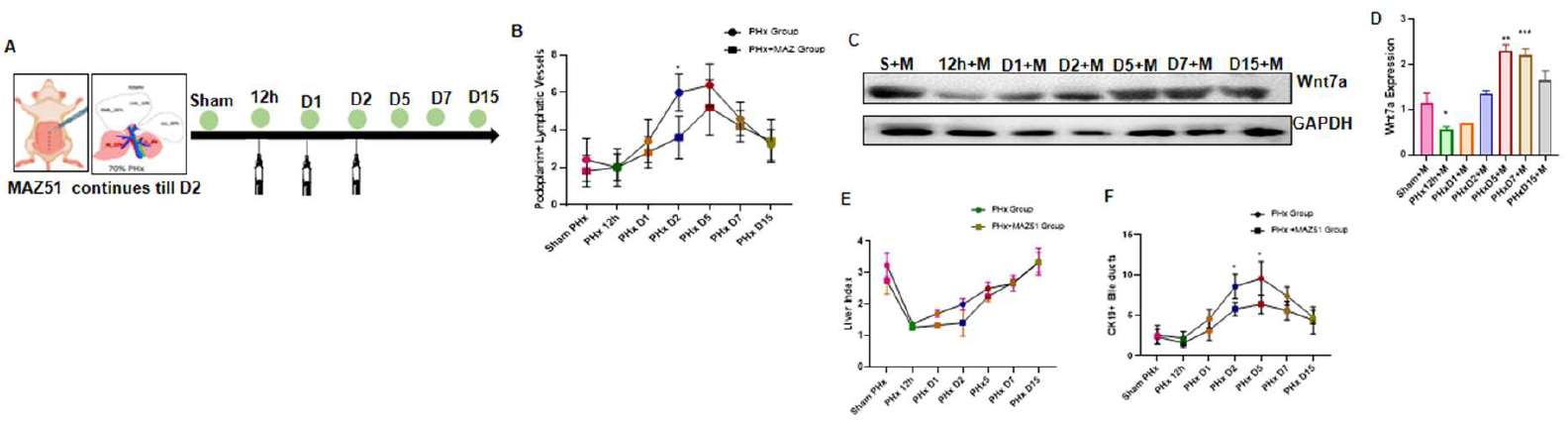
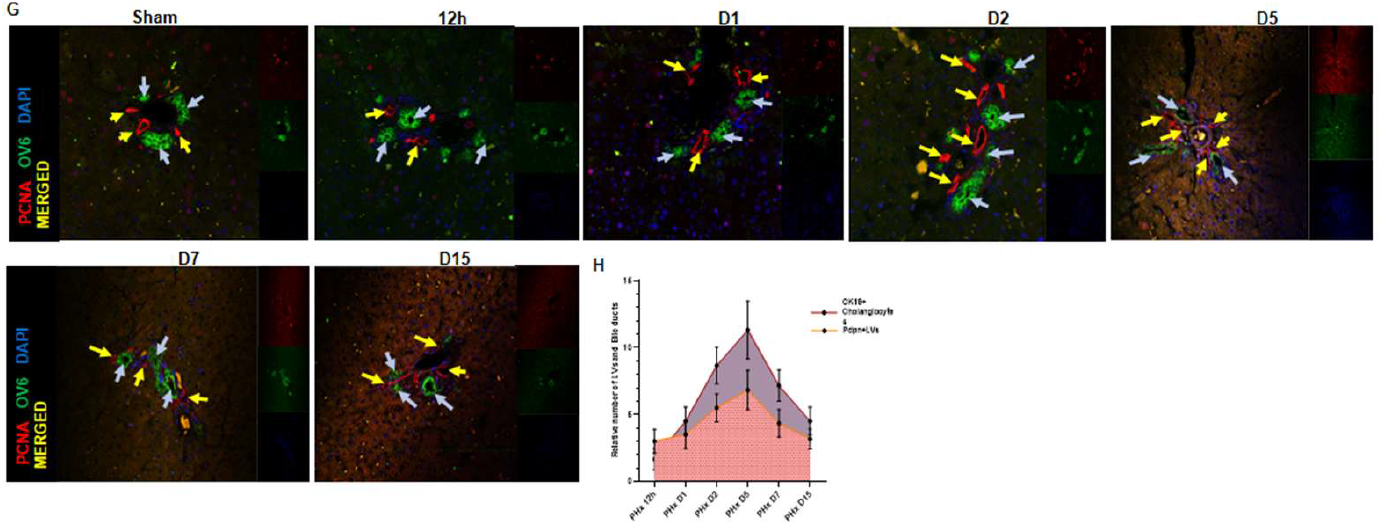
Schematic showing the time course and administration of MAZ51 (a VEGFR−3 inhibitor) in PHx models compared to the vehicle group. MAZ51 was administered intraperitoneally daily from the time of surgery until D2 post-PHx. Animal models were analyzed from 12h to D15 (**Fig4A**). Representative flow cytometry plots and quantification of LyECs (CD45−CD31+PDPN+) at D2 and D5 in MAZ51-treated PHx animals compared to the vehicle group (**Fig4B** n=3, *p <0.05). Representative Western blot analysis (**Fig4C**) and quantification of Wnt7a expression at different time point’s post-PHx in the MAZ51 group (**Fig4D**, *p <0.01, **p <0.01, ***p <0.001). Liver index (liver weight/body weight ratio) analysis in the MAZ51 PHx group compared to the vehicle at time points post PHx (**Fig4E**). Flow cytometry analysis and quantification of CK19+ cholangiocytes post-PHx in the MAZ51 group compared to the vehicle (**Fig4F**, *p <0.01). Representative dual immunofluorescence images and quantification showing the kinetic relationship between PDPN+ LVs (red) and CK19+ bile ducts (green) from 12h to D15 post PHx. Nuclei are stained with DAPI (blue) (**Fig4G&H**, n=3, mean + SD).

### Wnt7a Mediates Cholangiocyte Proliferation via FZD7/AKT/JAK1/STAT3 Signaling Pathway

Next, we wanted to establish a definitive relationship between Wnt7a and cholangiocytes proliferation. We first dissected the possible receptors of Wnt7a in the liver using string pathway analysis (**Fig5A**). The two major receptors observed were Frizzled 7(FZD7) (**Fig5B**) present on cholangiocytes and FZD5 present on hepatocytes (**SFig5A**). At D2 post-PHx, among all liver cells, gene expression of FZD7 was upregulated in cholangiocytes (**SFig5B**). FZD7 IHC also showed its localization mainly on the cholangiocytes in the periportal areas (**SFig5C&D**). Protein levels of FZD7 was also maximum at D2 (52.1±2.2%, p =0.01) post-PHx compared to sham (**Fig 5C&D**). After MAZ51 treatment, there was a reduction in FZD7 expression at D1 (36.3±2.2%, p =0.02) and D2 (45.6±4.3%, p =0.02) (**SFig5E&F**), indicating a possible interaction of Wn7a-FZD7.

**Figure 5:**
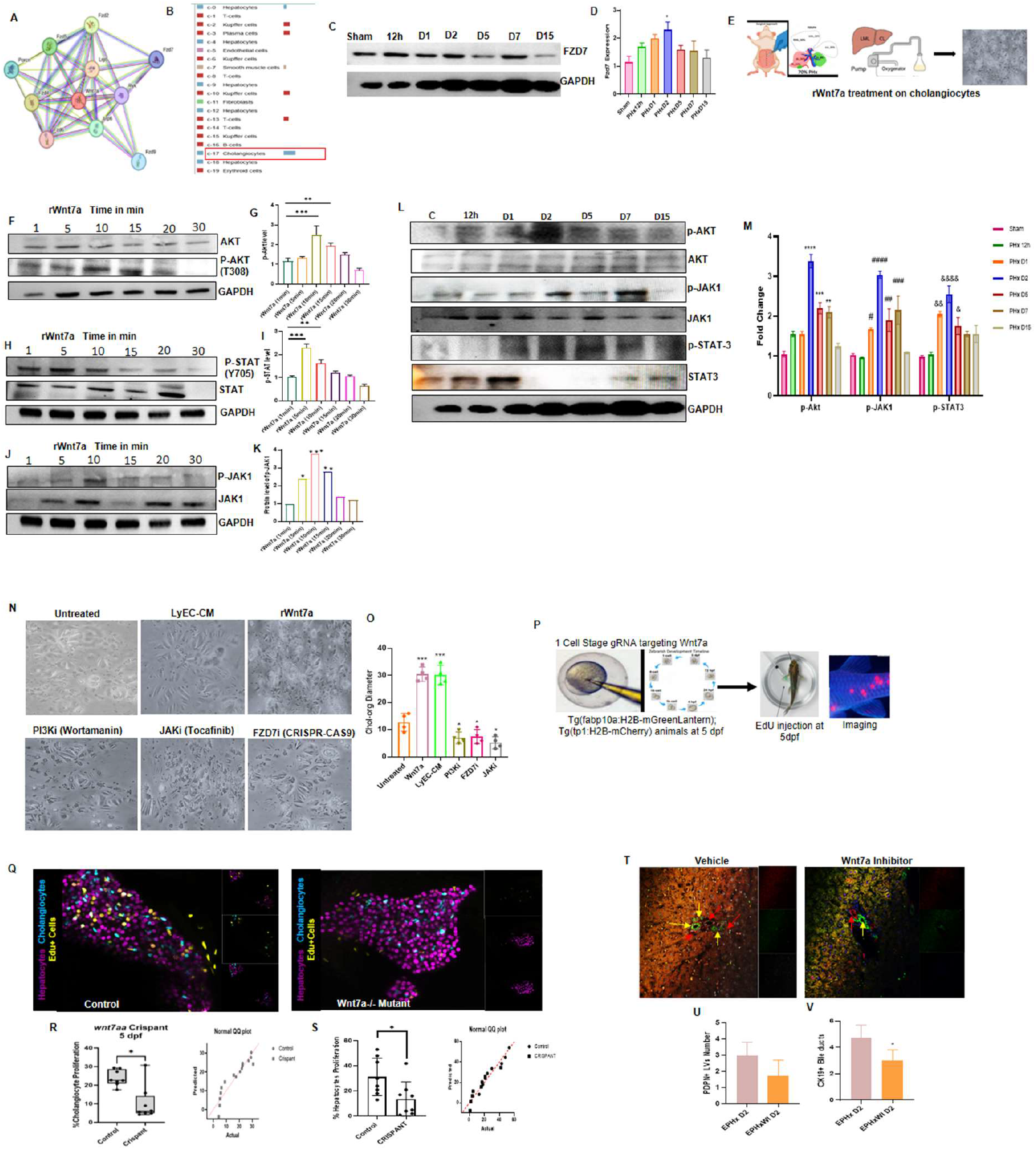
Wnt7a Signaling via the FZD7 Receptor Drives Cholangiocyte Proliferation and Liver Regeneration. Analysis using the STRING database identifying potential Wnt7a receptors involved in the activation of hepatic cells (**Fig5A**). Graphical representation indicating that FZD7 is a likely Wnt7a receptor expressed on the surface of cholangiocytes (**Fig5B**). Protein level analysis and quantification of FZD7 expression in isolated cholangiocytes from 12h to D15 post-PHx, showing maximum expression at D2 post-PHx (**Fig5C&D**, *p <0.01, **p <0.01). Schematic illustrating the time points for rWnt7a treatment of isolated cholangiocytes for molecular mechanism analysis (**Fig5E**). Protein level analysis and corresponding quantifications of isolated cholangiocytes treated with rWnt7a (**Fig4F-K**, *p <0.01, **p <0.01, ***p <0.001). Protein level analysis and quantification of p-AKT, p-STAT3, and p-JAK1 in rat cholangiocytes at various time points post-PHx (**Fig4L&M**, *p <0.01, **p <0.01, ***p <0.001). Representative images and quantification of 3D cholangiocyte organoid proliferation. Organoid growth is increased by LyEC-conditioned media (LyEC-CM) and recombinant Wnt7a (rWnt7a) but significantly reduced by the inhibitors Wortmannin (AKT inhibitor), Tocafinib (JAK/STAT3 inhibitor), and FZD7 gRNA (**Fig5N&O**, mean + SD, n=3, *p <0.01, ***p <0.001). EdU incorporation assay in Wnt7a Crispant (Wnt7a knockdown) zebrafish (Tg(fabp10a:H2B−mGreenLantern); Tg(tp1:H2B−mCherry)) compared to sham controls at 5 days post-fertilization (dpf) (**Fig5P**). Representative images show hepatocytes (purple), cholangiocytes (cyan), and EdU+ proliferating cells (yellow) (**Fig5J**). Quantification shows a significant reduction in EdU+ cholangiocytes ((**Fig5R, n=8**, mean + SD, *p <0.01) and EdU+ hepatocytes ((**Fig5S, n=8**, mean + SD, *p <0.01) in the Wnt7a Crispant model. Administration of Suramin (a Wnt7a inhibitor) via tail vein injection in 70% PHx rats results in a significant reduction in the number of CK19+ cholangiocytes and PDPN+ LVs compared to vehicle-treated controls, as shown by representative IHC images (**Fig5T-V**, n=3, mean + SD, *p <0.01).

Next, we studied the mechanisms underlying the growth-promoting effects of Wnt7a-FZD7 in the cholangiocytes. Wnt7a primarily regulates cell behaviour through the canonical Wnt/β-catenin pathway and non-canonical pathways, including the planar cell polarity (PCP) pathway. We treated the cholangiocytes with rWnt7a in vitro and collected the whole cell protein from 1 min to 30 minutes (**Fig5E**). Activation of p-AKT (2.27 fold, p<0.0001, **Fig4F&F**), p-STAT3 (2.3 fold, p<0.0001, **Fig5H&I**), and p-JAK1 (1.87 fold, p=0.001, **Fig4J&K**) was observed within 10min, 5 min, and 10 min respectively in cholangiocytes. However, no activation of β-catenin was observed, suggesting that Wnt7a acts via the PCP pathway in cholangiocytes. Next, we checked the in vivo activation of the PCP pathway in the cholangiocytes isolated from PHx models at different time points. We observed a clear activation of p-JAK1 (3.4 fold, p <0.0001), p-STAT3 (3.1 fold, p <0.0001) and p-AKT (2.7 fold, p <0.0001) maximally at D2 post-PHx compared to sham (**Fig4L&M**). In the MAZ51 treated group, the cholangiocytes showed a transient decrease in expression of downstream proteins, p-AKT (2.3 fold, p <0.0001), p-JAK1 (2.38 fold, p <0.001) and p-STAT3 (1.8 fold, p=0.03) at D2 post PHx as compared to untreated PHx animals (**SFig4G&H**). The expression of these proteins was normalized at D5 and D7. This clearly indicated that an inhibition of lymphangiogenesis that causes a decrease in Wnt7a reduced proliferation of cholangiocyte/bile ducts via inhibiting the PCP pathway.

To further validate the interaction between hepatic LyECs and cholangiocytes via Wnt7a-FZD7 pathway, we performed transfection studies using CRISPR-CAS9 based FZD7 inhibition in cholangiocytes. At plasmid concentration of 20-30ng/ml, out of these 3 plasmids synthesized, plasmid 3 was found to be most efficient for cell transfection studies, showing a 60% reduction in FZD7 expression in the cholangiocytes **(SFig5I&J, SFig5K&L**). FZD7 silencing, downstream pathway inhibition of the PCP pathway by Wortamanin (Akt inhibitor) and Tocafinib (JAK inhibitor) all showed similar effects of significant reduction of cholangiocyte proliferation by 75.0±1.98%, p=0.046, 54.5±12.7%, p=0.08 and 63.6±14.1%, p=0.014 respectively (**Fig5N&O)**.

Next, to assess how loss of WNT7A affects developmental proliferation of the liver, we performed in vivo CRISPR–Cas9 mutagenesis of the *wnt7aa* locus in zebrafish F0 crispants (**Fig5P**). One-cell stage embryos from *Tg(fabp10a:H2B-mGreenLantern); Tg(tp1:H2B-mCherry)* crosses were injected with a *wnt7aa*-targeting gRNA or a scramble control. At 5 dpf, proliferation was quantified by EdU pulse–chase (12 h) followed by imaging and cell-type–specific scoring in the double-transgenic background (**Fig5P**). Cholangiocyte proliferation was significantly lower in *Wnt7aa* crispants than in scramble-injected siblings, as visible in false-color renderings (cyan, cholangiocytes; yellow, EdU^+^ nuclei; **Fig5Q**). Quantitatively, *Wnt7aa* disruption produced a 2.34-fold, p=0.01 decrease in cholangiocyte proliferation (**Fig5R**). Hepatocyte proliferation was also reduced (2.57 fold, p=0.0162; **Fig5S**). Based on these results, in next set of 70% PHx rat models, we used an inhibitor of Wnt7a, i.e. suramin at a dose of 10mg/kg via tail vein to assess the specific effects of Wnt7a inhibition on liver regeneration. HE and MT of suramin-treated animals showed some injury response such as cellular infiltration, hepatocytes injury, portal vein congestion etc as compared to vehicle group (**SFig5M**). The liver index at D2 post PHx was decreased (3.5 fold, p <0.0001) in the inhibitor group compared to Wnt7a treated animals (**SFig5N**). PDPN+ LVs and CK19+ cholangiocytes were decreased in the suramin-treated group compared to vehicle (41.6±1.8%, p <0.09 and 43.6±3.4%, p=0.0319 respectively, **Fig5T-V**). The percentage of PCNA+ cells was also decreased in suramin treated animals as compared to vehicle, indicating an overall reduction in liver cell proliferation (26.3±4.5%, p <0.04, **SFig5O&P**). A reduced PAS positivity in the liver sections of inhibitor-treated animals as compared (53.06±3.2%, p =0.04) to vehicle was seen, suggesting a decrease in the number of functional hepatocytes (**SFig5Q&R**). Hence, our in vivo studies proved that Wnt7a is an important growth factor for proliferation of cholangiocytes during both liver developmental and regeneration in vivo.

### Wnt7a potentiates hepatocyte and cholangiocyte proliferation in extended hepatectomy/small size syndrome models

We next studied the effects of Wnt7a in small for size syndrome (SFSS) rat models. Here, we utilized 80% rat models of PHx as a model of SFSS. SFSS is a model representing a rapid loss of liver regeneration and its functions. The model exhibits several complications such as persistent cholestasis, coagulopathy, ascites, encephalopathy etc^19^. We utilized this model to test the efficacy of Wnt7a in promoting both hepatocyte and biliary proliferation. We injected rWnt7a via tail vein in SFSS models and compared them with SFSS vehicle animals at D2 post PHx (**Fig6A**). Liver index was significantly increased (6.6 fold, p <0.006), in the treated group compared to the vehicle groups (**Fig6B**). The liver architecture of treated SFSS models showed no significant changes in comparison to untreated PHx (**SFig6A&B**). Flow analysis showed an increased percentage of CK19+ cholangiocytes (60.6±2.2%, p <0.001) and CD31+PDPN+ LyECs (74.3±3.4%, p <0.001) in the treated animals as compared to the vehicle ones (**Fig6C&D**). The IF imaging for PDPN+ LVs and CK19+ cholangiocytes showed an enhanced number of both the cells in the treated group as compared to sham (53.6±5.2%, p =0.002, and 36.6±4.7, p=0.01, **6E-G**). An increased PAS+ area was observed in treated models in comparison to vehicle (52.4±6.4%, p=0.007, **6H&I**). An increased no. of PDPN+ LVs, CK19+ bile ducts and PCNA+ hepatocytes was also seen in treated groups versus the vehicle (52.3±3.8%, p =0.002, 36.6±3.5%, p=0.01, 29.4±4.5%, p=0.009 **SFig6C-F**). These results validated the growth-promoting effects of Wnt7a on both cholangiocytes and hepatocytes in vivo in rodent models. Next, we investigated if Wnt7a also mirrors these effects in human liver cell, specifically the cholangiocytes. Cholangiocytes isolated from the excised part of diseased human livers were cultured as chol-organoids and treated with rWnt7a for a period of one week with daily administration of rWnt7a (10-20ug/ml). There was a significant increase in the proliferation and formation of chol-organoids with enhanced PCNA positivity in the presence of rWnt7a compared to untreated controls (39.6±4.8%, p=0.006, **Fig6K&L**). The flow analysis also showed an increased proliferation rate. (**SFig 6G**). Next we tested the serum levels of Wnt7a in donors of LDLT at baseline, D1 & D2 and compared the Wnt7a secretion via ELISA. Details of these patients are shown in the table (7.**1**). Wnt7a serum levels were increased at D1 (15.17±3.1%, p=0.048) and more at D2 (16.19±3.4.2%, p =0.04) compared to the baseline (**Fig6J**). This suggested that Wnt7a levels can be detected early in human serum post-hepatectomy and it promotes the proliferation of human chol-organoids.

**Figure 6:**
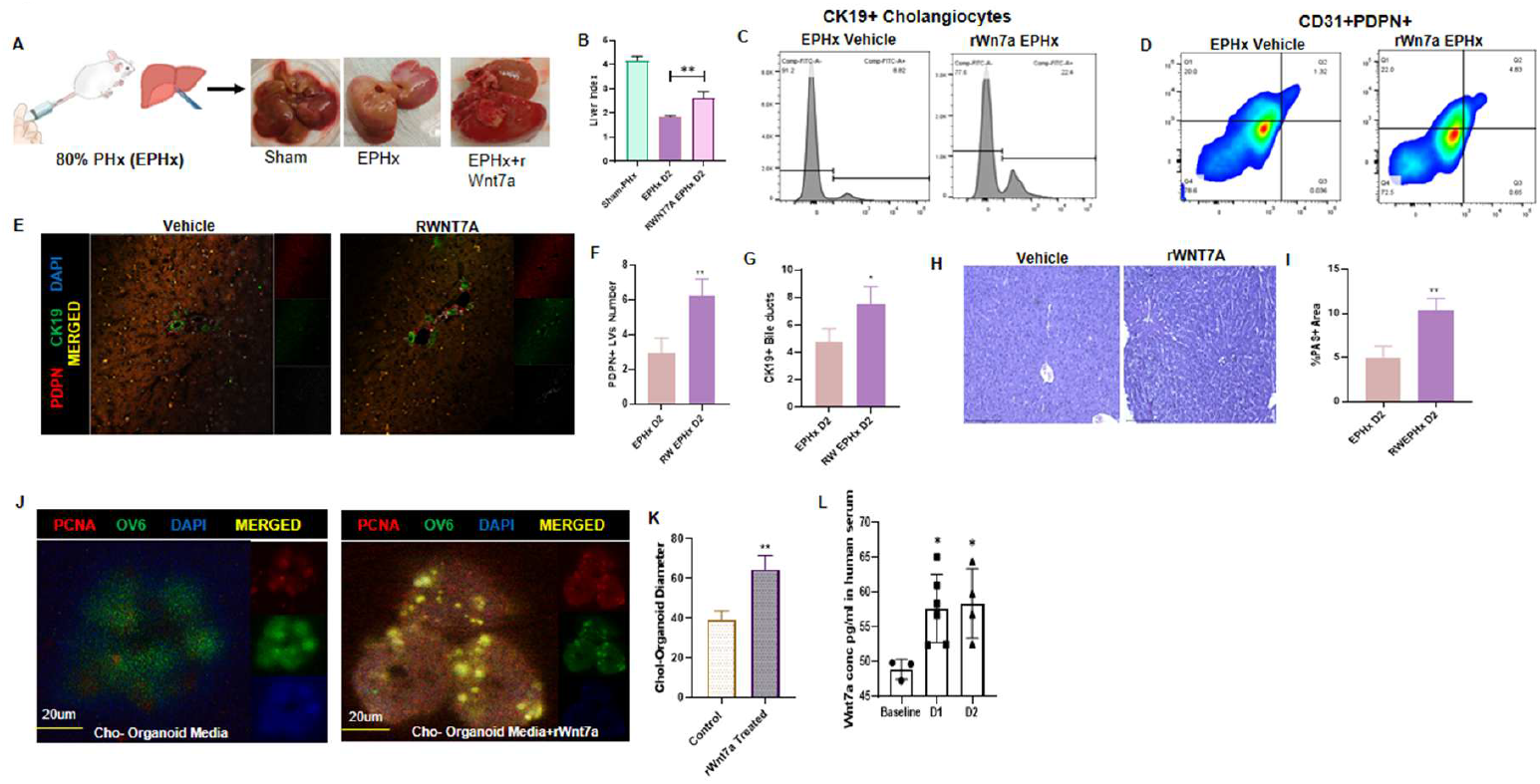
Therapeutic Administration of rWnt7a Enhances Regeneration in Models of Extended Hepatectomy (ePHx) and Small-for-Size Syndrome. Study plan for rWnt7a therapeutic intervention in 80% PHx (EPHx) models (**Fig6A**). Quantification of the Liver Index (liver weight/body weight ratio) at D2 post-EPHx, showing a significant increase in liver weight gain in rWnt7a-treated animals compared to vehicle controls (**Fig6B, n=3**, mean ± SD, *p <0.05). Flow cytometry analysis showing an increased number of CK19+ cholangiocytes (**Fig6C**) and CD45−CD31+PDPN+ LyECs (**Fig6D**) in rWnt7a-treated animals compared to vehicle controls post-EPHx. Representative immunofluorescence images and quantification of the number of bile ducts (BDs) and lymphatic vessels (LVs) (**Fig6E-G**, n=3, mean ± SD, *p <0.05, **p <0.01). PAS (Periodic acid– Schiff) staining images and quantification showing an increased number of PAS+ (glycogen-rich) hepatocytes in rWnt7a-treated EPHx animals (**Fig6H&I**, n=3, mean ± SD, **p <0.01). Representative images and quantification of cholangiocyte organoids isolated from explanted human recipient livers (**Fig6J&K**, n=3, mean ± SD, *p <0.05, **p <0.01). Wnt7a levels in human serum post-transplant surgery. ELISA analysis of Wnt7a levels in the serum of Living Donor Liver Transplant (LDLT) donors at baseline, post-PHx D1, and post-PHx D2 following surgery (**Fig6L**, n=6, mean ± SD, *p <0.05).

## Discussion

Liver regeneration after partial hepatectomy (PHx) is orchestrated by hepatocytes and cholangiocytes which are supported by angiocrine cues from non-parenchymal populations^20^. While hepatocyte proliferation is supported by multiple redundant angiocrine pathways, our data indicate that cholangiocyte growth depends critically on LyEC-derived Wnt ligand, Wnt7a. The contribution of hepatic lymphatic endothelial cells (LyECs) during liver regeneration remains poorly defined owing to technical challenges in their isolation and identification. Consistent with earlier reports that hepatic lymphatic vessels (LVs) are largely restricted to the portal triad^11^,^21^, we observed a marked expansion of LyECs around portal tracts during regeneration, peaking at day 5 post-PHx. Proteomic and functional analysis revealed Wnt7a as a key lymphangiocrine factor significantly upregulated at early time points post-PHx.

Functionally, recombinant Wnt7a promoted the proliferation of both hepatocytes and cholangiocytes in vitro. The zebrafish and mammalian models reinforced this observation, as genetic or pharmacological Wnt7a inhibition impaired both hepatocytes and cholangiocyte expansion, while Wnt7a supplementation in SFSS models enhanced biliary and hepatocyte proliferation. Interestingly, pharmacological inhibition of lymphangiogenesis with VEGFR3 inhibitor, MAZ51 that reduced LyEC numbers and Wnt7a expression, led to a marked decline in CK19+ cholangiocytes but not hepatocytes. This selective biliary effect highlights Wnt7a as a critical driver of cholangiocyte proliferation, while hepatocytes likely rely on redundant angiocrine signals from LSECs such as HGF^22^, Angpt2^23^, and Wnt2/9b^8^ after PHx. Despite these findings, we also acknowledge important limitations of our study. In zebrafish models, we could not definitively confirm LyECs as the cellular source of Wnt7a. Since zebrafish lack hepatic lymphatics during embryogenesis and only develop them around 5 dpf, Wnt7a in these contexts likely originates from other hepatic populations. To delineate the specific contribution of LyEC-derived Wnt7a, LyEC-specific Wnt7a knockout mice models will be valuable. Also, the reduction in CK19+ cholangiocytes upon VEGFR3 inhibition may be attributed to the concomitant loss of macrophages, since macrophages^24^ are key regulators of both lymphangiogenesis and ductular reaction. The precise interplay between LyECs, macrophages, and cholangiocytes therefore needs to be elaborated further. Nonetheless, our findings in rat PHx models and the spatial proximity of LyECs with portal tract cholangiocytes clearly support a niche-specific LyEC–Wnt7a axis in biliary regeneration.

Cholangiocyte proliferation post PHx has been less studied in comparison to hepatocyte proliferation in PHx models. The Hippo–YAP axis have been reported to regulate biliary cell fate decisions^24^,^25^, while the Notch pathway has been shown to be a principal driver of cholangiocyte proliferation following PHx^26^. Inhibition of Notch signaling reduces hepatic progenitor cell (HPC) differentiation into cholangiocytes, and disruption of Notch–RBPJ impairs bile duct regeneration by altering both proliferation and lineage specification^26^. Crosstalk between Notch and Hippo effectors, particularly RBPJ and YAP, is critical to maintain proper biliary architecture, with impairment of either pathway leading to abnormal biliary development and regeneration^26^,^27^. Our findings now add a new layer to this regulatory network by identifying LyEC-derived Wnt7a as a lymphangiocrine factor that sustains cholangiocyte proliferation during liver regeneration. Wnt7a acts via FZD7 to activate non-canonical JAK1/STAT3 and AKT pathways, distinct from canonical β-catenin signaling^28^. This is in concordance with recent single-cell RNA-seq studies showing that ductular Wnt signaling during biliary injury occur independently of LGR5/Axin2 activation^29^, and also with some cancer studies where Wnt7a–FZD7–STAT3 is known to promote self-renewal and survival in non-cancer models^30^,^31^,^28^. While earlier biliary injury models such as DDC or bile duct ligation (BDL) identified cholangiocytes as a source of Wnt7a driving ductular reactions^32^, our findings now establish LyECs as an additional, previously unrecognized source of Wnt7a during physiological liver regeneration. In our proteomics analysis of LyECs, we also observed an upregulation of other proteins/pathways in LyECs involved in cell proliferation such as PI3K, MAPK signalling pathways related to pluripotency of stem cells, hepatobiliary cells proliferation etc, re-enforcing an important but unrecognized role of LyECs in liver cell proliferation.

The translational relevance of Wnt7a was very well observed in the SFSS models, which mimics clinical post-hepatectomy liver failure. Studies have shown both compromised hepatocyte functions and endothelial fitness in the SFSS models and that a combined with G-CSF and HNF4α significantly improved functional liver recovery as well as the survival of the hepatectomized animals^33^. In this setting, Wnt7a treatment restored hepatocyte, vascular, and biliary compartments and improved regeneration, paralleling combinatorial strategies using hepatic growth factors and G-CSF. Furthermore, human cholangiocyte organoids responded robustly to recombinant Wnt7a, underscoring its therapeutic potential, especially in biliary regeneration. Moreover, elevated serum Wnt7a levels in human transplant donors post-PHx highlight its clinical relevance, although the direct contribution of LyEC-derived Wnt7a to bile duct regeneration in human settings remains to be fully defined.

Interestingly, Wnt7a exposure also induced expression of biliary markers in hepatocytes in vitro. This is in agreement with previous reports of Wnt-driven hepatocyte-to-cholangiocyte transdifferentiation in an injured liver^32^. Although we did not study this process in PHx models, yet such conversion is likely minimal in our models since both hepatocytes and cholangiocytes are capable of maintaining their own populations post PHx. The potential contribution of LyEC-derived Wnt7a to hepatocyte to cholangiocyte conversion remains an intriguing possibility that needs further exploration in biliary injury models. It is to be noted, however that an early upregulation of FZD5 in hepatocytes post-PHx in our experiments, do suggest that hepatocytes may indeed respond to Wnt7a through distinct receptor–ligand interactions.

In summary, we reveal that Wnt7a secreted by LyECs promotes both hepatocyte and cholangiocyte proliferation during early phases of liver regeneration, extending its role beyond injury-driven biliary repair. Our study underscores the importance of focusing on the role of these “rare” cell types in liver regeneration. The therapeutic effects of rWnt7a in SFSS models and in human cholangiocyte organoids, emphasizes its translational potential, particularly for improving liver functions in post-hepatectomy insufficiency and repairing of damaged bile ducts in cholangiopathies.

## Acknowledgement

SPS is supported by Fonds de la Recherche Scientifique (FNRS) grants 40027730 (CDR) and 40020360 (PDR) and Ramalingaswami Re-entry Fellowship from Department of Biotechnology (DBT).

