## supplementary for "Wnt7a is a Novel Lymphangiocrine Factor Driving Cholangiocyte Proliferation during Liver Regeneration": Supplementary Wnt7a Paper.pdf

### Supplementary Figures

Supplementary Figure 1

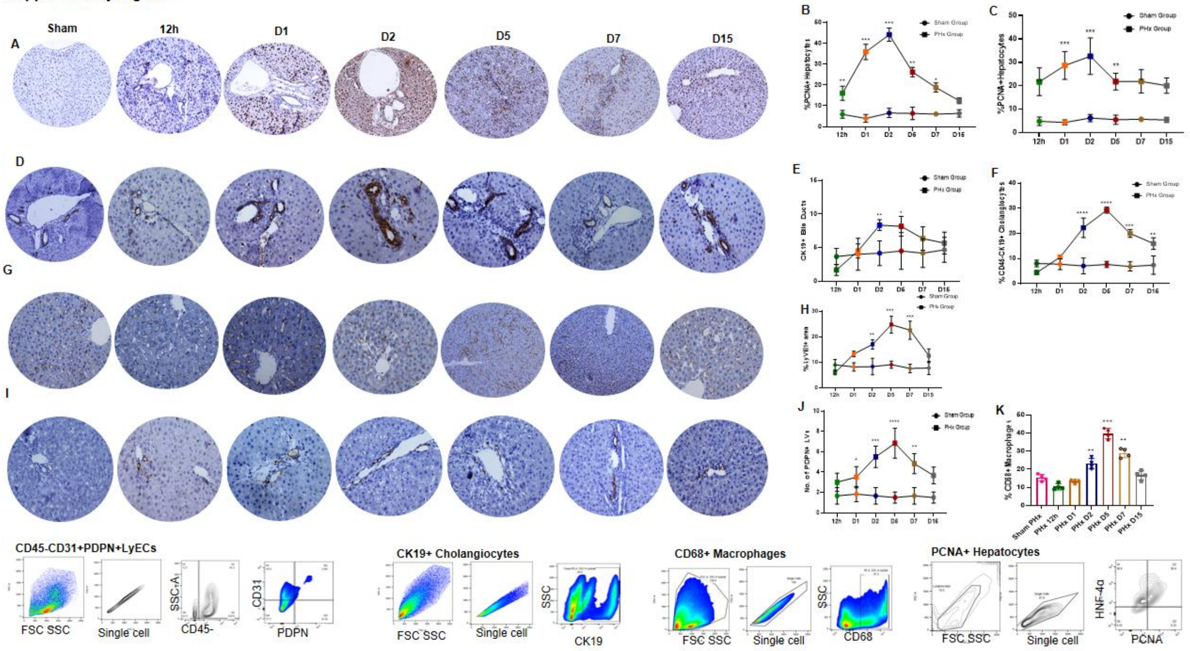


***Supplementary Figure 1:*** *PCNA showed an increased expression in liver tissues at D1 &D2 post-PHx (**SFig1A&B,** scale bar= 200µm,* mean ± SD, *n=3**,*** *p <0.01, **p <0.01, ***p <0.001*). Increased HNF4α+Ki67+ hepatocytes are shown in flow analysis (**SFig1C**). Maximum CK19+ cholangiocytes are shown in **SFig1D&E,** scale bar= 100µm*, mean ± SD, *n=6**,*** *p <0.01, **p <0.01, ***p <0.001*. Flow analysis data also supports the similar results **(SFig1F,** n=4**,*** mean ± SD, **p <0.01, ***p <0.001, ****p <0.0001***)**. IHC for CD31+ Endothelial cells also showed an increased expression mainly at D2&D5 (**SFig1G&H,** scale bar= 200µm* mean ± SD, *n=4**,*** *p <0.01, **p <0.01, ***p <0.001*). Podoplanin IHC data confirming LVs increased at D2 and D5 is shown in (**SFig1I&J,** scale bar= 100µm*, mean ± SD, *n=6**,*** *p <0.01, **p <0.01, ***p <0.001, ****p <0.0001*). Flow analysis of CD68+ macrophages found an increased no. at D2 post-PHx compared to sham (**SFig1K**,* mean ± SD, *n=4**,*** ***p <0.001, ****p <0.0001*). Flow analysis panel for LyECs, cholangiocytes, PCNA+ hepatocytes and macrophages (**SFig 1L**).*

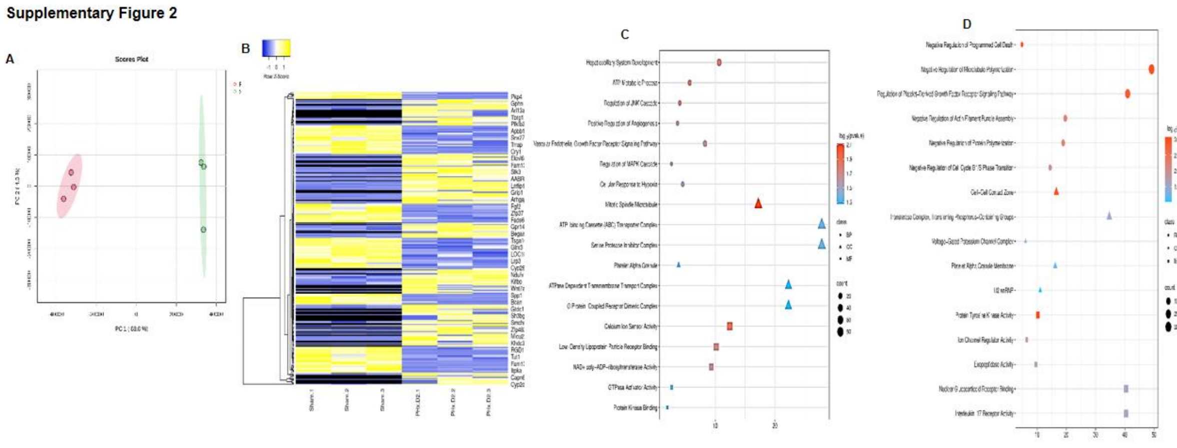


***Supplementary Figure 2:*** *PCA plot illustrating the separation and variation between LyECs from Sham and PHx D2 rats* ***(SFig2A,*** *n=3 in each group****).*** *Heatmap analysis showing the expression profiles of top 100 most significantly up-regulated and 100 most significantly down-regulated genes between Sham LyECs and PHx D2 LyECs* ***(SFig2B,*** *n=3 in each group****).*** *Gene Ontology (GO) terms showing significantly up-regulated and down-regulated in LyECs at PHx D2 compared to Sham, categorized by biological process, cellular component, and molecular function* ***(SFig2C&D).***

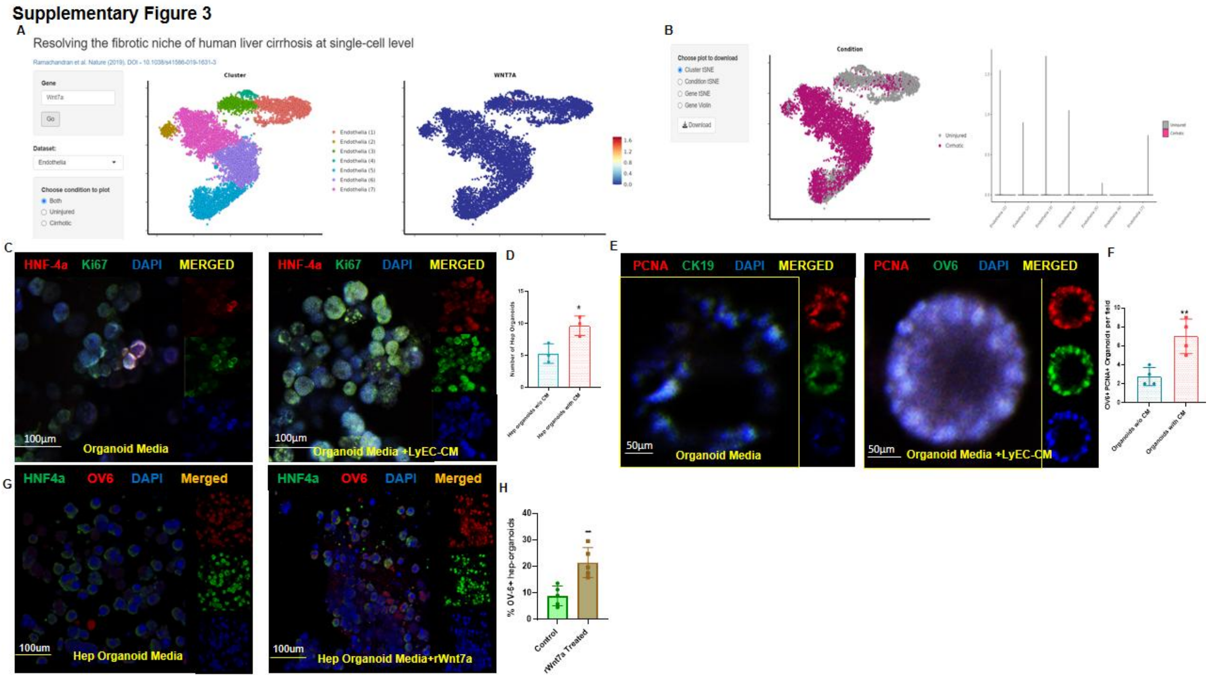


***Supplementary Figure 3-*** *Analysis of existing databases showing the source of Wnt7a expression originating from two distinct endothelial cell populations: Liver Sinusoidal Endothelial Cells (LSECs) and LyECs in human fibrotic livers (****SFig3A&B****). Representative immunofluorescence images and quantification of hepatocyte organoid diameter when cultured with LyEC-CM compared to control. Nuclei are stained with DAPI (blue), HNF−4α (hepatocyte marker, red), and Ki67 (proliferation marker, green) (****SFig3C&D,*** *p <0.05*). Representative immunofluorescence images and quantification of*

*diameter showing a 2.23-fold increase in cholangiocyte organoid diameter when cultured with LyEC-CM compared to control. Nuclei are stained with DAPI (blue), PCNA (proliferation marker, red), and CK19 (cholangiocytes marker, green) (**SFig3E&F,** *p <0.05*). Immunofluorescence imaging of hepatocytes cultured with rWnt7a. Nuclei are stained with DAPI (blue), HNF−4α (hepatocyte marker, red), and OV−6 (biliary marker, green) (**SFig3G&H,** *p <0.05*).*

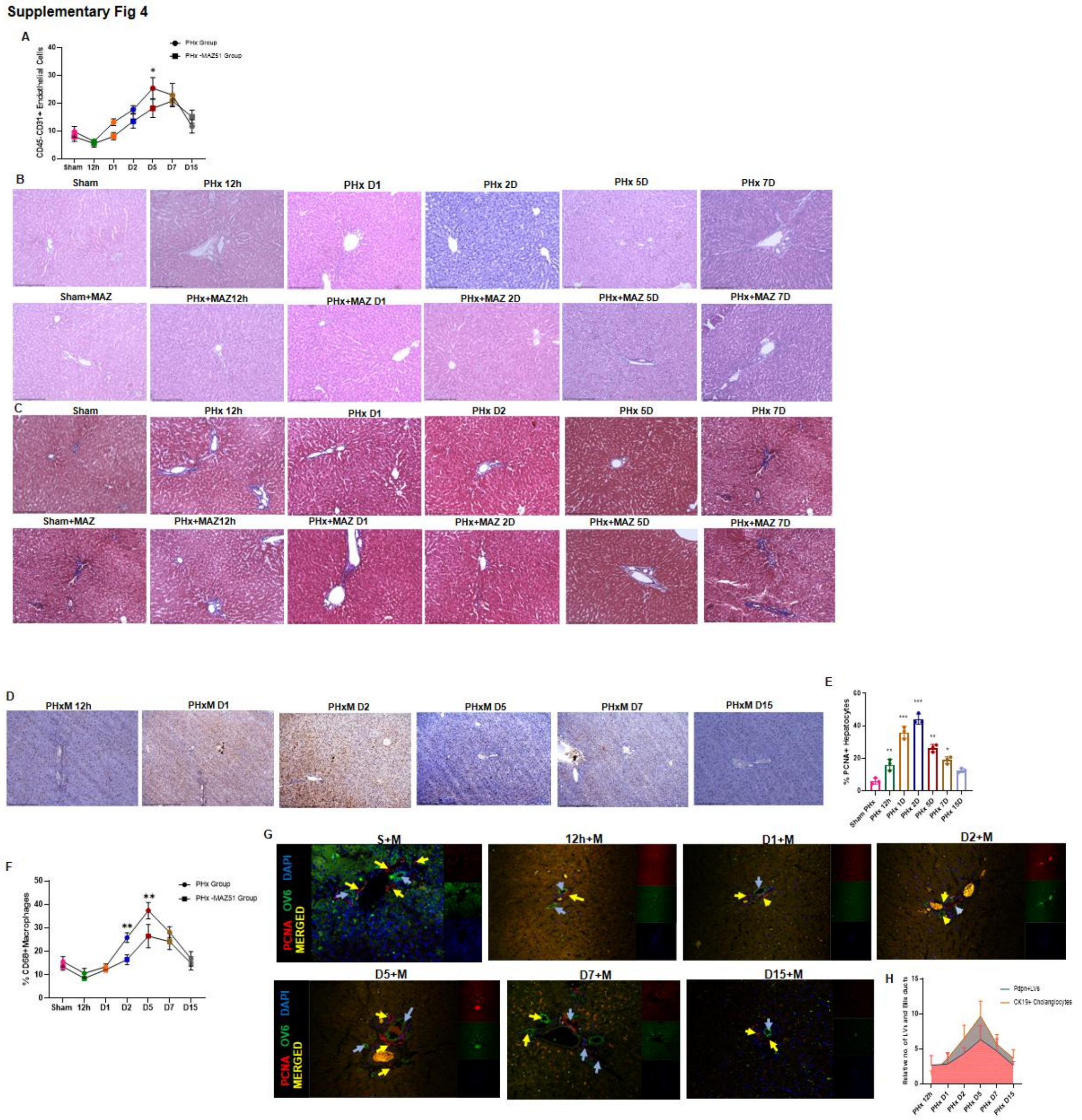


***Supplementary Figure 4***

*Flow cytometry analysis of LECs (CD45−CD31+) in the PHx MAZ51 group compared to the PHx vehicle group (**SFig4A,** *p <0.01*). Representative histology images of PHx MAZ51 compared to PHx* *vehicle. Scale bars: 200µm (**SFig4B&C**). Immunohistochemistry (IHC) for PCNA+ cells in PHx and PHx MAZ51 groups (**SFig4D&E,** n=3, mean ± SD,* *p <0.01, **p <0.01*). Flow analysis for CD68+ macrophages at D2 post-PHx in the MAZ51 treated group compared to vehicle (**SFig4F**). Dual immunofluorescence imaging and quantification comparing the kinetics of PDPN+ LVs (red) and CK19+ bile ducts (green) from 12h to D15 in PHx MAZ51 compared to the PHx vehicle group. Nuclei are stained with DAPI (blue) (**SFig4G&H,** n=3, mean ± SD,* *p <0.01, **p <0.01*).*

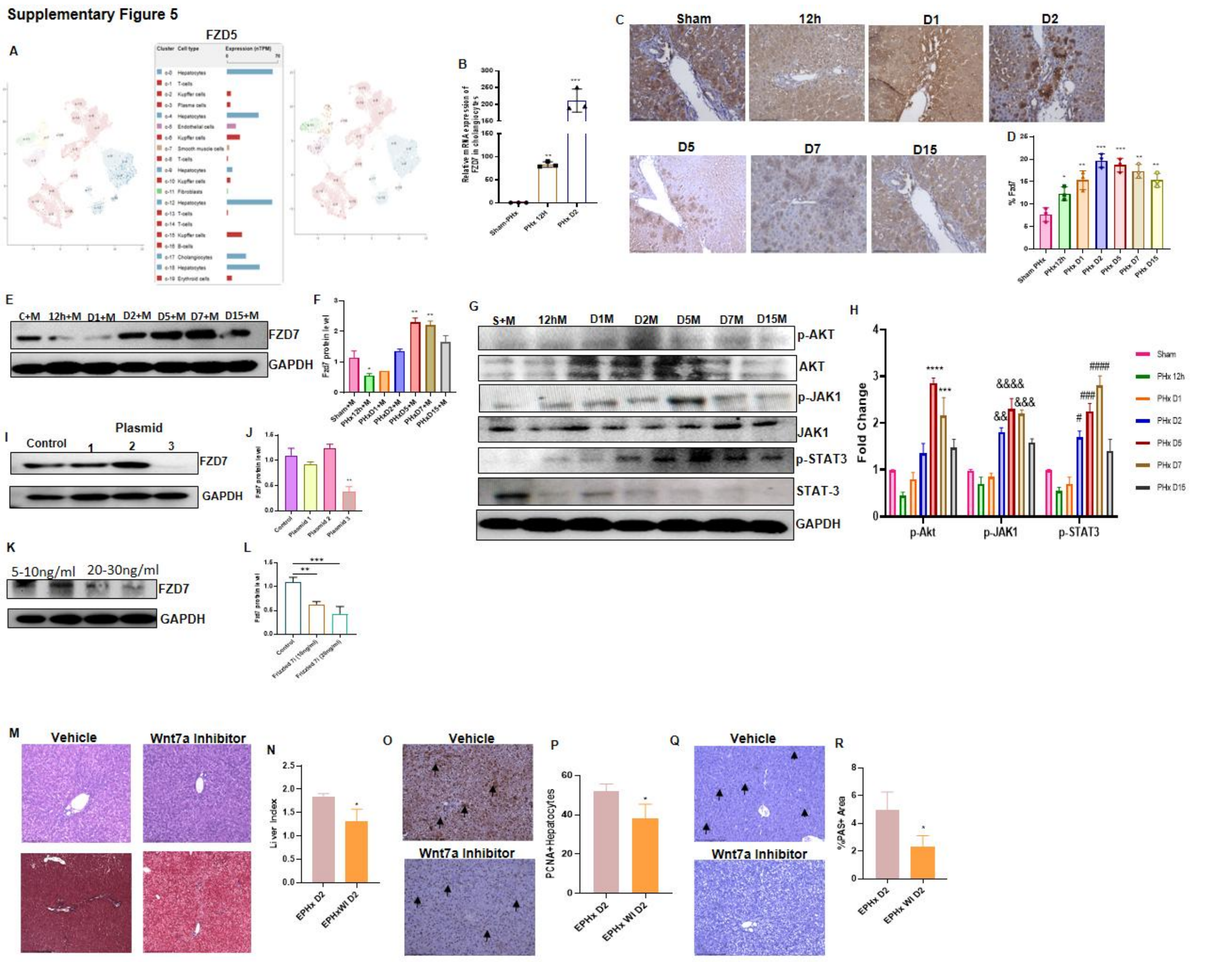


***Supplementary Figure 5***

*Analysis of Protein Cell Atlas database identifying FZD5 as another potential Wnt7a receptor expressed on the surface of hepatocytes **(SFig5A)**. Quantitative RT-PCR analysis showing increased FZD5 mRNA expression in isolated hepatocytes at D2 post-PHx compared to Sham (**SFig5B, n=3,** mean ± SD,* **p <0.01, ***p <0.01*). Representative Immunohistochemistry (IHC) images **(SFig5C)** and quantification confirming the increased expression of FZD7 in the liver at D2 post-PHx compared to Sham (**SFig5D, n=3,** mean ± SD,* *p <0.05, **p <0.01, ***p <0.01*). Protein level analysis and quantification of FZD7 in PHx MAZ51 treated animals, showing a delayed expression with peaks at D5 and D7 compared to the D2 peak observed in the vehicle group (**SFig5E&F,** *p <0.05, **p <0.01). ). Western blot analysis and quantification of p-AKT, p-STAT3, and p-JAK1 in isolated cholangiocytes* *from D2 to D5 and D7 post-PHx (**SFig4G&H,*** *p <0.01, **p <0.01*). Representative data and quantification showing that FZD7 gRNA plasmid 3 demonstrated the highest transfection efficiency in cultured cells compared to other plasmid constructs (**SFig5I&J,*** *p <0.05*). Representative data and quantification of FZD7 receptor loss in cultured cholangiocytes following transfection with FZD7 gRNA plasmid (20-30 ng/mL concentration), demonstrating approximately 60% receptor knockdown **(Fig5K&L,*** **p <0.01, ***p <0.001***). Histological analysis (e.g., H&E) showing increased evidence of post-operative injury in Suramin-treated PHx models compared to vehicle, including increased portal vein congestion, cellular infiltration, and hepatocyte injury (**SFig5M**). Analysis of the Liver Index (liver weight/body weight ratio) showing a significant reduction in liver mass in Suramin-treated models compared to vehicle controls following PHx (**SFig5N, n=3,** mean ± SD,* *p <0.05*). Representative IHC images and quantification of PCNA+ cells showing a significant reduction in the proliferation of hepatic cells (hepatocytes and potentially others) in Suramin-treated PHx models compared to vehicles (**SFig5O&P,** n=3, mean ± SD,* *p <0.05. *PAS (Periodic acid–Schiff) staining for glycogen, indicating functional status of hepatocytes, shows a reduced number of PAS+ hepatocytes in Suramin-treated models compared to vehicle, suggesting impaired hepatocyte function (**SFig5Q&R**, n=3, mean ± SD,* *p <0.05*).*

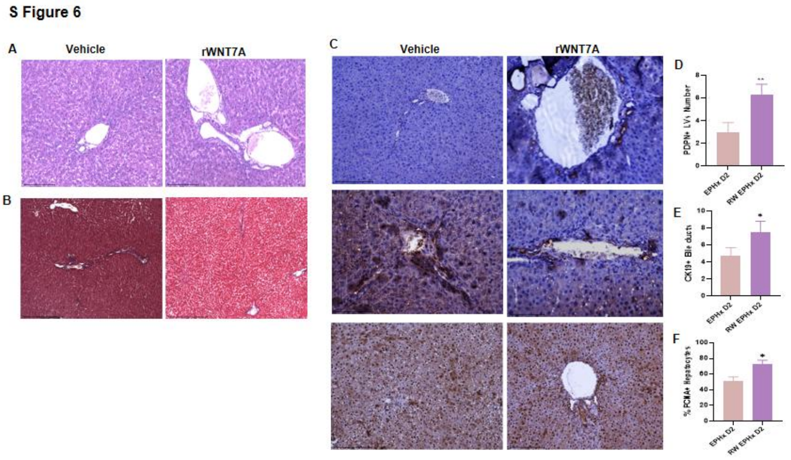


***Supplementary Figure 6: Histological and Immunohistochemical Analysis of the Liver Following rWnt7a Treatment in EPHx Models.***

*Representative images of H&E staining and Masson's Trichrome (MT) staining comparing rWnt7a-treated and vehicle-treated 80% PHx (EPHx) livers (**SFig6A&B**). (C) Representative images of immunohistochemistry (IHC) performed on liver tissue sections from both rWnt7a and vehicle groups for the quantification of PDPN+ LVs, CK19+ bile ducts, and PCNA+ proliferating hepatic cells (**SFig6C**). Quantification of IHC data showing a significant increase in the number of PDPN+ Lymphatic Vessels (LVs) in the rWnt7a-treated group compared to vehicle control (**SFig6D,** n=3, mean ± SD,* *p <0.05, **p <0.01*). Quantification of IHC data showing a significant increase in the number of CK19+ bile ducts in the rWnt7a-treated group compared to vehicle control (**SFig6E, n=3,** mean ± SD,* *p <0.05, **p <0.01*). Quantification of IHC data showing a significant increase in the number of PCNA+ proliferating hepatic* _cells in the rWnt7a-treated group compared to vehicle control (**SFig6F,** n=3, mean_ ± _SD,_ *p <0.05, **p <0.01_)._
